# Neuronal responses support a role for orbitofrontal cortex in cognitive set reconfiguration

**DOI:** 10.1101/082610

**Authors:** Brianna J. Sleezer, Giuliana Loconte, Meghan D. Castagno, Benjamin Y. Hayden

**Affiliations:** Department of Brain and Cognitive Sciences, Center for Visual Science and Center for the Origins of Cognition University of Rochester

**Author notes:** Corresponding author: Benjamin Y. Hayden, Department of Brain and Cognitive Sciences, University of Rochester, Rochester, NY 14618.

**Keywords:** switching, executive control, rule, wisconsin card sorting task, rhesus macaque

## Abstract

We are often faced with the need to abandon no-longer beneficial rules and adopt new ones. This process, known as cognitive set reconfiguration, is a hallmark of executive control. Although cognitive functions like reconfiguration are most often associated with dorsal prefrontal structures, recent evidence suggests that the orbitofrontal cortex (OFC) may play an important role as well. We recorded activity of OFC neurons while rhesus macaques performed a version of the Wisconsin Card Sorting Task that involved a trial-and-error stage. OFC neurons demonstrated two types of switch-related activity, an early (switch-away) signal and a late (switch-to) signal, when the new task set was established. We also found a pattern of *match modulation*: a significant change in activity for the stimulus that matched the current rule (and would therefore be selected). These results extend our understanding of the executive functions of the OFC. They also allow us to directly compare OFC with complementary datasets we previously collected in ventral (VS) and dorsal (DS) striatum. Although both effects are observed in all three areas, the timing of responses aligns OFC more closely with DS than with VS.

## INTRODUCTION

The orbitofrontal cortex (OFC) is a critical site for decision-making and adaptive behavior. Its contributions to the evaluation and comparison of rewards are well established (Padoa-Schioppa and Assad, 2006; Wallis, 2007). Perhaps less well-known are its executive roles. OFC is critical for linking stimuli to values, in monitoring consequences of actions, in detecting and resolving conflict, in metacognition, and encoding rules and storing sensory information in working memory (Abe and Lee, 2011; Kepecs et al., 2008; Lara et al., 2009; Mansouri et al., 2014; Rushworth et al., 2011; Schoenbaum et al., 1999; Sleezer et al., 2016; Strait et al., 2016; Tsujimoto et al., 2009; Wallis and Miller, 2003; Wallis et al., 2001). Indeed, the list of executive functions of the OFC is almost as long as those associated with classical executive structures like dorsolateral prefrontal cortex (DLPFC) and dorsal anterior cingulate cortex (dACC).

One executive function for which the role of OFC is not as well understood is cognitive set reconfiguration. This term refers to the adjustment of cognitive strategies or mental representations in response to changing goals or environmental circumstances (Robbins, 2007). It is often called switching for short. A classic example of switching is recognizing that a familiar driving route to work is blocked and identifying and changing to an alternative route. Switch-related signaling is a classic executive function and is most closely associated with executive regions in the dorsal prefrontal cortex and parietal cortex (Alan et al., 1994; Dias et al., 1996a; Kamigaki et al., 2012; Mansouri et al., 2006). Specific evidence for this linkage comes, in part, from physiological studies showing systematic modulations of firing rate during switch trials relative to other trials.

Although a number of studies suggest that OFC contributes to simpler types of flexible decision-making (Dias et al., 1996a, 1996b, 1997; McAlonan and Brown, 2003), there are several reasons to believe that OFC may participate directly in switching, especially between rules. First, as noted above, recent evidence supports a role for OFC in many executive functions. Second, we and others have delineated a role for OFC in maintenance of rules or in updating based on rules (Buckley et al., 2009; Sleezer et al., 2016; Tsujimoto et al., 2011; Wallis et al., 2001; Yamada et al., 2010). Third, recent work indicates that OFC lesions disrupt the ability to switch between behavioral rules in rodents (Birrell and Brown, 2000). Finally, we previously showed switching-related activity in two basal ganglia structures - the dorsal (DS) and ventral (VS) striatum-which are not generally associated with switching. Specifically, we found that switch signals in VS were strongest when switching away from previously relevant rules, while switch signals in DS were strongest when switching toward newly relevant rules. Because OFC has direct anatomical projections to both VS and DS, we hypothesized that it may have a direct role in switching as well.

In our earlier study on striatal contributions to switching, we also found that the appearance of switch signals in VS and DS was consistent with the appearance of associative learning signals (i.e. the systematic enhancement or suppression in firing rate for task-relevant targets when they appear in a sequence of stimuli) in both regions. We therefore wondered whether OFC neurons would demonstrate similar patterns of activity. Such signals likely relate to executive function, as they correspond to a linkage between mental representations of rules and presentation of specific offers.

To examine the role of OFC in switching, we recorded the activity of single neurons as macaques performed a version of the Wisconsin Card Sorting Task. In our version, rules were never cued, so subjects had to go through a trial-and-error phase to determine the currently relevant rule (this normally took 3-4 trials). They could then take advantage of the newly learned rule and maintain responding until the rule changed again (blocks were 15 trials long). We found systematic changes in firing associated with both early (switch-away) and late (switch-to) switch trials - i.e. explicit switch signals - in OFC neurons. We also found that switch signals in OFC were consistent with the appearance of associative learning signals in this region; associative signals arose slowly and only became strong once the rule was established. Our results are consistent with the idea that switch signals are linked to associative learning, and may even serve to initiate learning processes during flexible rule updating. More generally, these findings endorse a broader executive role for OFC and are consistent with a recent theory proposing that OFC instantiates a cognitive map of task space (Schuck et al., 2016; Wilson et al., 2014).

## MATERIALS AND METHODS

### Surgical procedures

All animal procedures were approved by the University Committee on Animal Resources at the University of Rochester and were designed and conducted in compliance with the Public Health Service’s Guide for the Care and Use of Animals. Two male rhesus macaques *(Macaca mulatta)* served as subjects. We used standard electrophysiological techniques as described previously (Strait et al., 2014). A small prosthesis for holding the head was used. Animals were habituated to laboratory conditions and then trained to perform oculomotor tasks for liquid reward. A Cilux recording chamber (Crist Instruments) was placed over the OFC. Position was verified by magnetic resonance imaging with the aid of a Brainsight system (Rogue Research Inc.). Animals received appropriate analgesics and antibiotics after all procedures. Recording locations care shown in **Figure 1C**. Throughout both behavioral and physiological recording sessions, the chamber was kept sterile with regular antibiotic washes and sealed with sterile caps.

**Figure 1.**
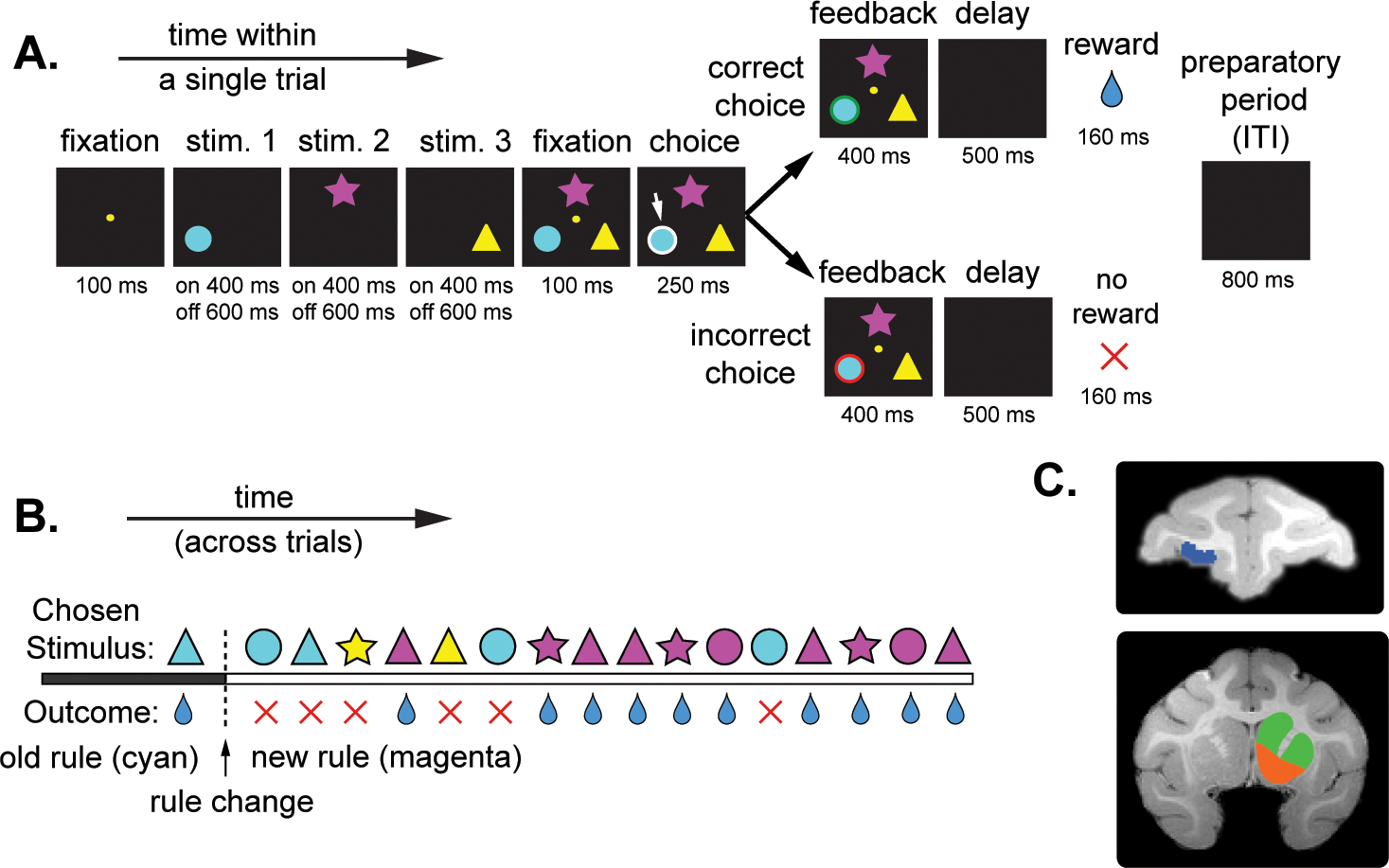
Task and recording locations. (A) Timeline of Wisconsin Card Sorting Task. (B) Example block. In this example, the correct rule is magenta. (C) Magnetic resonance image of subject C. Recordings were made in orbitofrontal cortex (highlighted in blue). Comparison sites were in the ventral striatum (highlighted in orange), and dorsal striatum (highlighted in green).

### Recording sites

We approached OFC through a standard recording grid (Crist Instruments). We used the standard atlas for all area definitions (Paxinos et al., 2000). We defined OFC as the coronal planes situated between 29 and 36 mm rostral to the interaural plane, the horizontal planes situated between 0 and 9 mm from the ventral surface, and lateral to the medial orbital sulcus. We recorded from Area 13m (Öngür and Price, 2000). We confirmed recording locations before each recording session using our Brainsight system with structural magnetic resonance images taken before the experiment. Neuroimaging was performed at the Rochester Center for Brain Imaging, on a Siemens 3T MAGNETOM Trio Tim using 0.5 mm voxels. We confirmed recording locations by listening for characteristic sounds of white and gray matter during recording, which in all cases matched the loci indicated by the Brainsight system. The Brainsight system typically offers an error of <1 mm in the horizontal plane and <2 mm in the z-direction.

### Electrophysiological techniques

Single electrodes (Frederick Haer & Co., impedance range 0.8 to 4M Ω) were lowered using a microdrive (NAN Instruments) until waveforms between 1 and 3 neuron(s) were isolated. Individual action potentials were isolated on a Plexon system (Plexon Inc., Dallas, TX). Neurons were selected for study solely on the basis of the quality of isolation; we never pre-selected based on task-related response properties.

### Eye-tracking and reward delivery

Eye position was sampled at 1000 Hz by an infrared eye-monitoring camera system (SR Research). Stimuli were controlled by a computer running Matlab (Mathworks) with Psychtoolbox and Eyelink Toolbox. Visual stimuli were presented on a computer monitor placed 57 cm from the animal and centered on its eyes. A standard solenoid valve controlled the duration of juice delivery. The relationship between solenoid open time and juice volume was established and confirmed before, during, and after recording.

### Behavioral task

The task described here is the same as that used in two previous manuscripts (Sleezer and Hayden, 2016; Sleezer et al., 2016). Subjects performed an implementation of the Wisconsin Card Sorting Task (WCST, Moore et al., 2005). Our version of the task uses two dimensions (color and shape) and six specific rules (three shapes: circle, star, and triangle, and three colors: cyan, magenta, and yellow, **Figure 1A**). On each trial, three stimuli were presented asynchronously at the top, bottom left, and bottom right of the screen (1 second asynchrony). The color, shape, position, and order of stimuli were fully randomized on each trial. Each stimulus was presented for 400 ms and was followed by a 600 ms blank period. Subjects were free to fixate upon the stimuli when they appeared. Then all three stimuli reappeared simultaneously with a central fixation spot. The subject fixated on the central spot for 100 ms and then indicated its choice by shifting gaze to its preferred stimulus and maintaining fixation on it for 250 ms. Failure to maintain gaze for 250 ms did not lead to the end of the trial, but instead returned the subject to a choice state; thus, subjects were free to change their mind if they did so within 250 ms (although in our observations, they almost never did so). Following a successful 250 ms fixation, visual feedback was provided (a green/red outline around the chosen stimulus for correct/incorrect choices, respectively). After visual feedback, there was a 500 ms blank delay period; correct choices were followed by a liquid (water) reward. All trials were separated by an 800 ms inter-trial interval (ITI), which we refer to as the preparatory period. In each block, subjects responded according to one of the six rules. Subjects were required to use a trial-and-error learning process to determine the correct rule. Rule changes occurred after 15 correct trials and were not explicitly cued.

### Analysis of behavioral performance across different types of rule changes

To determine whether subjects’ performance differed depending on the type of rule change that occurred at the beginning of the block, we calculated the average number of trials monkeys completed prior to rule acquisition (i.e. the first trial in a series of four consecutive correct trials, Sleezer and Hayden, 2016) following intra-dimensional and extra-dimensional rule changes. Intra-dimensional rule changes refer to instances when the rule change occurs within one rule category (i.e. color to color or shape to shape), while extra-dimensional rule changes refer to instances when the rule change occurs across rule categories (i.e. color to shape or shape to color). To compare the number of trials monkeys completed prior to rule acquisition across intra-dimensional and extra-dimensional switches, we used a two-way repeated measures ANOVA with the between subjects factor subject (Monkey B, Monkey C) and the within subjects factor block type (intra-dimensional, extra-dimensional). We used post-hoc Fisher’s LSD tests to compare specific differences across groups.

### Analysis of switch-related neural activity

On the first trial of each block, subjects almost always chose according to the previously relevant rule. Because the block transition was not explicitly cued, we called this the “inevitable error trial”. On blocks where the new rule happened to match the previous one by chance (1/6 of blocks), the first trial did not produce an error, and monkeys did not change strategy, so, for purposes of analysis, we treated these blocks as 30 trial blocks. Moreover, because there were three stimuli on each trial, with two dimensions each, occasionally (1/3 of blocks), the correct stimulus on the first trial was consistent with the previously relevant rule. We therefore specified in our definition of the inevitable error trial that it referred to the first trial on which choosing according to the previous rule would produce an error.

We examined switch-related neural activity during the 1,420 ms post-feedback period following feedback (that is, the combined duration of the delay, reward, and preparatory periods) and prior to the start of switch and non-switch trials. We analyzed this period because monkeys likely reconfigured their cognitive rule set on switch trials during this period. Non-switch trials were defined as all trials other than switch trials. The two types of switch trials are defined below.

We identified two points in the block when monkeys likely reconfigured (i.e. switched) their cognitive rule set. The ***early switch*** was the post-feedback period following an incorrect choice and immediately prior to the start of the first correct trial. We chose this trial because subjects had switched but had not yet begun consistently responding according to the new rule. We excluded early switch points that were also identified as late switch points. The ***late switch point*** was on the trial immediately following the first trial of at least 4 consecutive correct trials.

Task-related activity during the post-feedback period was determined using ANOVA with the factors trial type (switch or nonswitch), block type (intradimensional or extra-dimensional), trial outcome (reward or no-reward), and next trial outcome (reward or no-reward). In these analyses, trial outcome refers to the outcome during the reward period during the post-feedback period, while next trial outcome refers to the outcome during the reward period on the following trial. Although we were interested in the effects of trial type, block type, and their interaction, we included trial outcome and next trial outcome in the ANOVA model to control for the potential influence of reward or error related activity. Because current trial outcome and next trial outcome were not fully crossed with trial type in this model (that is, switch trials always consisted of a non-rewarded trial followed by a rewarded trial), we used a nested ANOVA in which current and next trial outcome were nested in trial type. A nested ANOVA measures the effects of a factor while partialling out the effects of a nesting factor. Thus, by utilizing a nested ANOVA in which current and next trial outcome were nested in trial type, this model includes an estimate of the effects of current and next trial outcome, which thus serves as control for reward outcome related effects. We conducted these analyses separately for early and late switch points. Based on the ANOVA results, we classified task-related activity into two types. The first type showed a significant main effect (*P* < 0.05) of trial type, the second showed a significant interaction (*P* < 0.05) between trial type and block type. Post-hoc comparisons (Fisher’s LSD test) were conducted if the interaction was significant (*P* < 0.05). We refer to neurons with a main effect of trial type as *general switch signaling neurons* and neurons with an interaction between trial type and block type as *context-specific switch signaling neurons.*

To determine if the proportion of cells demonstrating a significant switch-related effect (a main effect of trial type or a significant interaction between trial type and block type) was significantly above chance, we conducted binomial tests, and adjusted the p-value using a Bonferroni correction for two comparisons. We corrected for two comparisons because we analyzed activity at both early and late switch points. We chose to maintain an alpha of 0.05 and multiply the resultant p-values by two as a way of implementing the Bonferroni correction. Thus, the p-values reported for binomial tests in this paper have been adjusted for two comparisons, where appropriate. To determine if proportions of cells demonstrating an effect were significantly different across OFC, VS and DS, we implemented a mixed model binary logistic regression procedure using the between subjects factor brain region (OFC, VS, DS) and the within subjects factors trial period (early, late) and modulation type (general switch, context dependent switch). The model was fit using a generalized estimating equation (GEE) procedure, implemented in SPSS. In this analysis, “within subjects” and “between subjects” refer to neurons. In this procedure, an omnibus Wald Chi-Square test was applied to determine the significance of group effects, followed by pairwise comparisons using Fisher’s LSD tests to examine specific group effects.

To examine the percent of variance explained by each switch-related effect across the populations of OFC, VS and DS neurons, we calculated the average partial η^2^. Partial η^2^ is a measure of effect size in ANOVA, which measures the proportion of variance attributable to a factor after partialling out other factors from the non-error variance (Cohen, 1973). Partial η^2^ is calculated as:

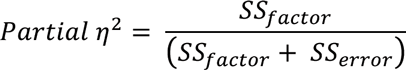

where SS_factor_ is the variation attributable to the factor (sum of squares for the factor), and SSerror is the error variation (sum of squares error). To compare the average partial η^2^ for switch-related effects at early and late switch points and in OFC, VS and DS, we used a two-way repeated measures ANOVA with the factors brain region (OFC, VS and DS) and switch period (early and late), followed by post-hoc Fisher’s LSD tests.

### Analysis of Associative Learning-Related Activity

To examine associative learning-related neural activity, we calculated the average firing rate during each of the three stimulus presentation epochs on all correct trials. We defined the stimulus presentation epoch as the 1000 ms period consisting of 400 ms when the stimulus was on the screen and the following 600 ms when the stimulus was off the screen. We then used two-way t-tests to compare the average firing rate during epochs in which the correct stimulus was presented to the average firing rate during epochs in which the correct stimulus was not presented.

To examine the magnitude of correct stimulus selectivity, we calculated Hedge’s g, a measure of effect size similar to Cohen’s d. Hedge’s g is recommended when groups have different sizes, and was also developed to remove a positive bias affecting Cohen’s d (Hedges, 1981). Since the sample sizes for the presentation of incorrect stimuli were always larger than the sample sizes for the presentation of correct stimuli (since each trial consisted of one correct stimulus and two incorrect stimuli), we chose to calculate effect size using Hedge’s g, rather than Cohen’s d. Hedge’s g is calculated as:

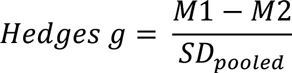

where M1 and M2 are the means of each group, and SD_pooled_ is the pooled standard deviation, calculated as:

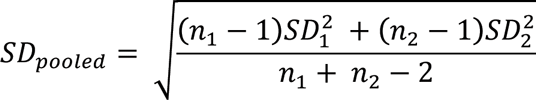

where n_1_ and n_2_ are the sample sizes for each group, and SD_1_ and SD_2_ are the standard deviations for each group.

To compare selectivity across brain regions, we first determined the average time of maximum selectivity within trials in each region (averaged across all correct trials) and analyzed a 200 ms period surrounding that time (100 ms before and 100 ms after). We then used these analysis epochs to compare selectivity in OFC, VS, and DS before and after late switch points. We compared selectivity across switch periods and brain regions using a 2-way repeated-measures ANOVA with the factors switch period (pre-late switch and post-late switch) and brain region (OFC, VS, and DS), following by post-hoc Fisher’s LSD tests.

Statistical analyses were carried out using MATLAB release 2012b (MathWorks Inc), SPSS Statistics version 24 (IBM Analytics), and GraphPad Prism version 6 (GraphPad Software).

## RESULTS

### Behavioral Performance

After a 2-3 month period of training, both subjects were able to reliably learn new rules and maintain a high level of accuracy once new rules were acquired (**Figure 2A**). Once training was complete, we collected data during 78 individual OFC recording sessions (n = 35 sessions for subject B and n = 43 sessions for subject C). Subjects completed an average of 565.09 ± 12.57 (mean ± sem) trials per session and an average of 30.92 ± 0.83 blocks per session.

**Figure 2.**
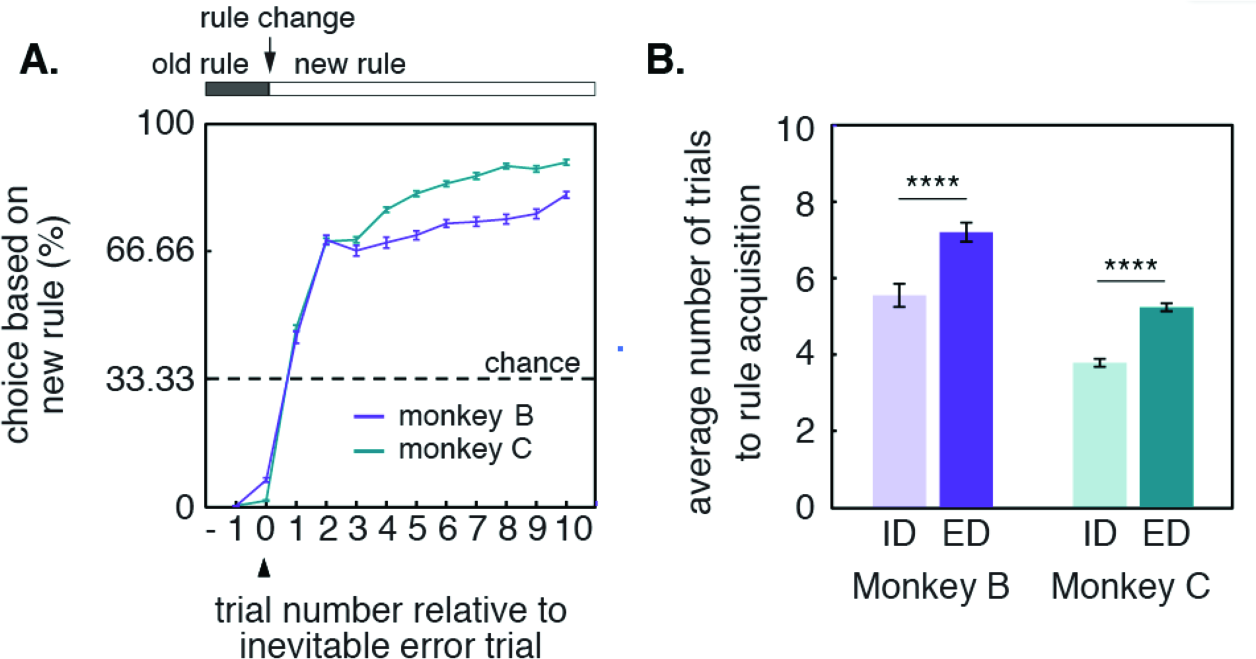
Behavioral performance. (A) Average choice accuracy on trials surrounding rule changes. (B) Average number of trials to rule acquisition following intra-dimensional (ID) and extra-dimensional (ED) rule changes.

On each block (besides the first block of each session), subjects completed either an intra-dimensional (ID) or extra-dimensional (ED) switch to the new rule. To determine whether subjects’ performance differed following intra-dimensional and extra-dimensional rule changes, we examined the number of trials monkeys completed prior to rule acquisition following intra-dimensional and extra-dimensional rule changes using a two-way repeated measures ANOVA with the between subjects factor subject (Monkey B, Monkey C) and the within subjects factor block type (intra-dimensional, extradimensional). This analysis revealed a significant main effect of subject (*F*(1,267) = 75.70; *P* < 0.0001) and of block type (*F*(1,267) = 136.2; *P* < 0.0001), but no interaction between the two (*F*(1,267) = 0.7783; *P* = 0.3785). The results of our post-hoc comparisons are shown in **Figure 2B**. We found that both subjects completed more trials prior to rule acquisition following extra-dimensional rule changes compared to intra-dimensional rule changes (**Figure 2B**, *P* < 0.0001 for both subjects, Fisher’s LSD Tests).

On average, subjects completed 14.94 ± 5.79 early switches and 27.62 ± 9.53 late switches per session. Prior to early switch trials, monkeys completed 3.12 ± 0.66 trials (3.34 ± 0.77 for monkey B and 2.97 ± 0.52 for monkey C), and prior to late switch trials, monkeys completed an average of 6.69 ± 2.31 (8.04 ± 2.74 for monkey B and 5.74 ± 1.29 for monkey C). These numbers include the inevitable error trial.

### Neurons in the orbitofrontal cortex demonstrate switch-related activity

We first characterized neural responses associated with switch trials. To do this, we compared firing rates on non-switch trials (all trials besides early- and late-switch trials) with those obtained on early switch trials (that is, the first correct trial after a switch). Then, in a separate analysis, we compared non-switch trials with late switch trials (the first correct trial in a series of at least 4 consecutive correct trials). We analyzed firing rate activity during the post-feedback period separately for each cell using ANOVA (see Methods).

Our assessment of switch-related activity focused on both preponderance and effect size during the post-feedback period, measured by the proportion of cells demonstrating a significant effect and the proportion of variance explained (partial η^2^) by the main effect of trial type (“general switch” modulation) and the interaction between trial type and block type (“context-dependent switch” modulation).

**Figure 3A** shows an example of an OFC neuron demonstrating general switch-related activity at both early and late switch points. The average firing rate response for this neuron was significantly greater on early switch trials than non-switch switch trials for both ID switches (red line and black dotted line, *P* < 0.0001, Fisher’s LSD test) and ED switches (pink line and gray dotted line, *P* = 0.0015, Fisher’s LSD test), and also significantly greater on late switch trials than non-switch trials for ID switches (dark blue line and black dotted line, *P* < 0.0001, Fisher’s LSD test) and ED switches (light blue line and gray dotted line, *P* < 0.0001, Fisher’s LSD test). This neuron showed no response difference for ID versus ED switches, whether looking at early switch points (red line and pink line, *P* = 0.9263, Fisher’s LSD test), or late switch points (dark blue line and light blue line, *P* = 0.6720, Fisher’s LSD test).

**Figure 3.**
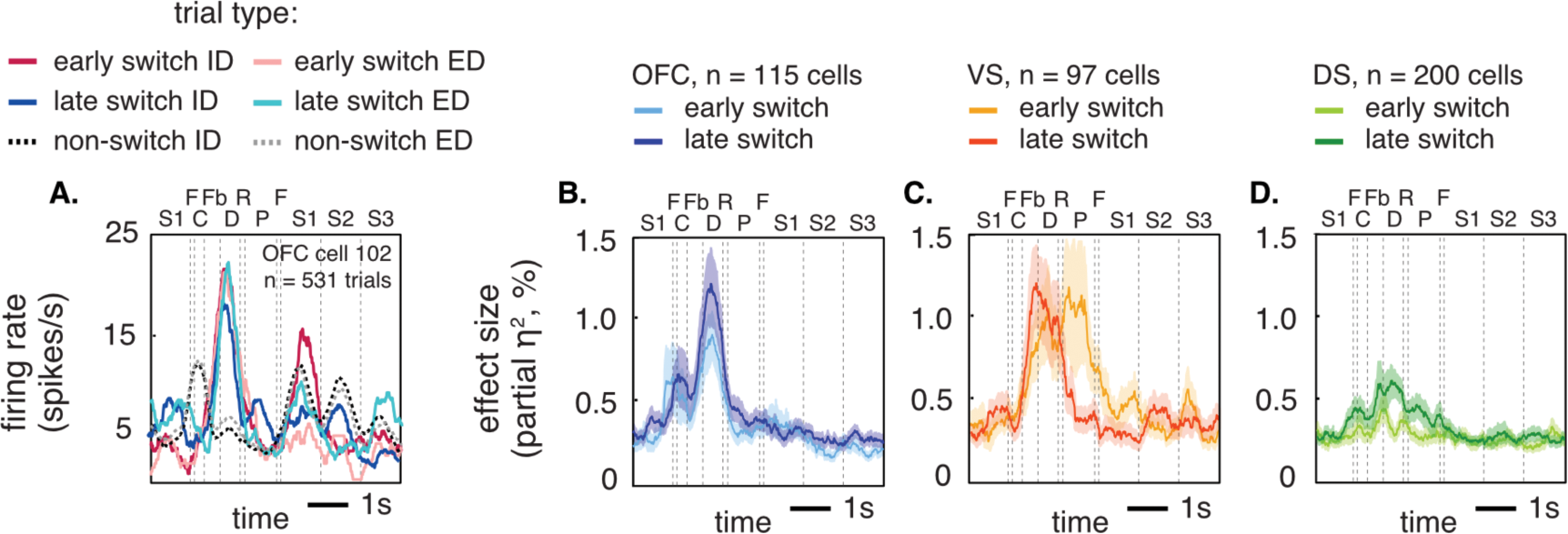
General switch modulation. (A) Average response of a single OFC neuron demonstrating general switch-related activity (i.e. a main effect of trial type [switch or non-switch]) at early and late switch points. Red and pink lines indicate intra-dimensional (ID) and extra-dimensional (ED) switch trials at early switch points. Blue and light blue lines indicate ID and ED switch trials at late switch points. Black and gray dotted lines indicate ID and ED non-switch trials. C, choice; Fb, feedback; D, delay; R, reward; P, preparatory period (ITI); F, fixation; S1, first stimulus appearance; S2, second stimulus appearance; S3, third stimulus appearance. (B-D) Proportion of variance explained (partial η^2^) across the populations of OFC (B), VS (C), and DS (D) neurons. Effect size measures reflect averages across all neurons (excluding 6 from VS and 4 from DS that were excluded due to an insufficient number of trials).

To determine whether the proportion of OFC cells demonstrating general switch modulation was above chance, we calculated the average proportion of cells demonstrating a significant effect across the 1,420 ms post-feedback epoch and performed binomial tests. We corrected for two comparisons (because we looked at early and late switch points) using a Bonferroni correction. These results are shown in **Figure 4A**. We found that the proportion of cells demonstrating general switch modulation was significantly above chance at early switch points (n = 20/115 cells) and at late switch points (n = 30/115 cells, *P* < 0.0001 in both cases).

**Figure 4.**
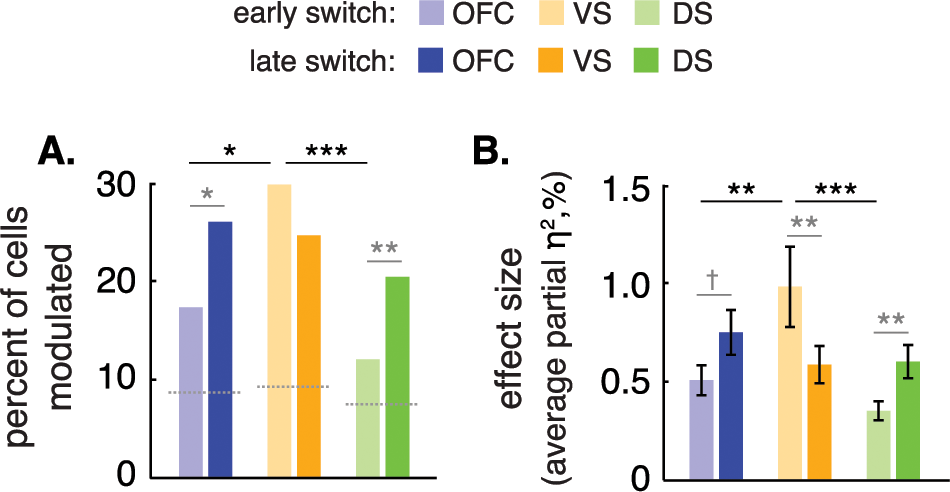
General switch modulation at early and late switch points. (A) Proportion of cells demonstrating a significant main effect of trial type at early and late switch points. (B) Proportion of variance explained (partial η^2^) by the main effect of trial type at early and late switch points. OFC (light blue and dark blue bars), VS (light orange and dark orange bars), and DS (light green and dark green lines). Bar graph shows the mean partial η^2^ (± SEM) during the post feedback period. Black lines and asterisks indicate significant effects across time (i.e. early and late switch points). Gray lines and asterisks indicate significant effects across brain regions. † *P* = 0.0502, * *P* < 0.05, ** *P* < 0.01, *\*\*\* P* < 0.001

In a previously published report, we examined the same effects in two striatal regions, ventral (VS) and dorsal (DS) striatum (Sleezer and Hayden, 2016). Data for VS and DS are shown as well, for comparison (**Figure 4A**). We found that the number of cells demonstrating switch modulation in these areas was significant at early switch points (VS: n = 29/97 cells; DS: n = 24/200 cells, Bonferroni adjusted *P* < 0.0001 for all three regions, corrected for two comparisons, binomial test) and at late switch points (VS: n = 24/97 cells; DS: n = 41/200 cells, Bonferroni adjusted *P* < 0.0001 for all three regions, corrected for two comparisons).

To compare the proportions of cells demonstrating each type of modulation across brain regions and across early and late switch points, we implemented a mixed model binary logistic regression procedure using the between subjects factor brain region (OFC, VS, DS) and the within subjects factors trial period (early, late) and modulation type (general switch, context dependent switch). In this analysis, the terms “within subjects” and “between subjects” refer to neurons. In this procedure, an omnibus Wald Chi-Square test was applied to determine the significance of group effects, followed by Fisher’s LSD tests to examine specific group differences. This analysis revealed a significant main effect of brain region (χ^2^ = 6.4922, *P* = 0.0389), a significant main effect of modulation type (χ^2^ = 68.3523, *P* < 0.0001), and a significant interaction between brain region and trail period (χ^2^ = 8.1057, *P* = 0.0174).

Pairwise comparisons are shown in **Figure 4A**. These analyses revealed a significantly greater proportion of cells demonstrating general switch modulation at late switch points compare to early switch points in both OFC and DS (OFC: *P* = 0.0461, DS: *P* = 0.0090, Fisher’s LSD tests). In contrast, we found no difference in the proportion of cells demonstrating general switch modulation at early and late switch points in VS *(P* = 0.3672). However, we did find that the proportion of cells demonstrating general switch modulation at early switch points was significantly greater in VS compared to both OFC and DS at the same time point (VS vs. OFC: *P* = 0.0322; VS vs. DS: *P* = 0.0006).

**Figure 3B** shows the average proportion of variance explained (partial η^2^) by the main effect of trial type in OFC across time within trials. **Figures 3C, and 3D** show the corresponding data in VS, and DS for comparison. To compare the average partial η^2^ across OFC, VS, and DS, we calculated the average partial η^2^ across the 1,420 sec post-feedback epoch and used a mixed-model ANOVA with the between subjects factors brain region (OFC, VS, and DS), and the within subjects factors trial period (early, late), and modulation type (general switch, context-dependent switch). As above, in this analysis, “within subjects” and “between subjects” refer to neurons. This analysis revealed a main effect of brain region (F_(2,409)_ = 4.6948, *P* = 0.0096), a main effect of modulation type (F_(1,409)_ = 62.4911, *P* < 0.0001), and an interaction between brain region and trial period (F_(2,409)_ = 9.9800, *P* = < 0.0001). Post-hoc comparisons are shown in **Figure 4B**.

We found a greater modulation in OFC at late switch points compared to early switch points, although this effect was not significant (*P* = 0.0502, Fisher’s LSD test). At early switch points, we found a greater magnitude of general switch modulation in VS compared to both OFC and DS (*P* = 0.0029 and *P* < 0.0001), but no difference between OFC and DS at the same time point (*P* = 0.2482). At late switch points, we found no difference between any of the three regions (OFC vs. VS: *P* = 0.3027; OFC vs. DS: *P* = 0.2694; VS vs. DS: *P* = 0.9174).

These results indicate that VS neurons demonstrate greater switch-related modulation during early periods of trial-and-error learning compared to later periods of rule acquisition and compared to OFC and DS neurons at the same time point. In addition, OFC, VS, and DS appear to demonstrate equal levels of switch-related modulation at later points of rule acquisition, while OFC and DS neurons demonstrate greater switch-related modulation during later periods of rule acquisition compared to early periods of trial-and-error learning.

### Latencies of switching signals

We next examined the latency of general switch signal appearance in OFC, VS, and DS at early and late switch points. None of these analyses were reported in our earlier study (Sleezer et al., 2016). To estimate latency, we calculated the average partial η^2^ for the main effect of trial type (switch or non-switch) across time within trials using a 50 ms sliding window slid in 10 ms steps across the 1,420 ms post-feedback period. To determine whether these latencies were significantly different across populations of neurons, we calculated the time of maximum selectivity across neurons and performed a one-way ANOVA using the factor brain region (OFC, VS, DS), separately at early and late switch points. This analysis revealed no significant differences in group latencies across brain regions at early *(P* = 0.1027) or late switch points *(P* = 0.5793).

### Context-specific switch signals arise during the early trial-and-error period in VS, but not OFC or DS

We next investigated context-specific switching activity (i.e. encoding of switches specific to either extra-or intra-dimensional switches, but not both). These results are similar to those reported in our previous paper (Sleezer and Hayden, 2016), however, the analysis technique used here is more sensitive and the OFC data were not reported in the previous paper. We found that the proportion of cells demonstrating context-dependent switch modulation at early switch points was significantly above chance in VS (n = 11/97 cells, Bonferroni adjusted *P* = 0.0067, corrected for two comparisons, binomial test), but not OFC or DS (OFC: n = 3/115 cells, Bonferroni adjusted *P* = 1.6642; DS: n = 9/200 cells, Bonferroni adjusted *P* = 1.0906, corrected for two comparisons). At late switch points, the proportion of cells demonstrating a significant effect was not significant in OFC, VS, or DS (OFC: n = 6/115 cells, Bonferroni adjusted *P* = 0.7051; VS: n = 5/97 cells, Bonferroni adjusted *P* = 0.7410; DS: n = 15/200 cells, Bonferroni adjusted *P* = 0.0887, corrected for two comparisons). In comparing proportions across brain regions at early switch points, we found a significantly greater proportion of cells demonstrating context-dependent switch modulation in VS compared to OFC (*P* = 0.0138) and a greater proportion of cells in VS compared to DS, which was marginally significant (*P* = 0.0542). We found no difference between OFC and DS at early switch points (*P* = 0.2033), or between any of the three regions at late switch points (OFC vs. VS: *P* = 0.9836; OFC vs. DS: *P* = 0.4128; VS vs. DS: *P* = 0.4241).

In comparing the strength of context-dependent switch modulation across early and late switch points, we found significantly greater modulation in VS at early switch points compared to late switch points (*P* = 0.0004, Fisher’s LSD test), but no difference between the two time points in OFC or DS (*P* = 0.7328, *P* = 0.3469). In comparing the strength of modulation across brain regions at early switch points, we found significantly greater modulation in VS compared to both OFC and DS (*P* < 0.0001 for both comparisons). We found no difference between OFC and DS at early switch points (*P* = 0.8194), nor did we find any differences between the three regions at late switch points (OFC: *P* = 0.9855, VS: *P* = 0.5200, DS: *P* = 0.5564).

Taken together with our findings regarding general switch-related activity, the above results suggest that VS neurons demonstrate greater switch modulation during the early trial-and-error period of the block compared to the later point of rule acquisition, and that a portion of these cells carry information about the rule context (i.e. whether the switch is intra-dimensional or extra-dimensional). In contrast, OFC and DS neurons demonstrate greater switch modulation during the later period of the block, and these signals carry no information regarding the switch context.

### Neurons in the OFC demonstrate associative learning related activity

We next wanted to know how OFC responses reflect learning of associations between stimuli and outcomes (i.e. reward or no reward), and how these responses relate to switch modulation. To do this, we examined the neural response to the three probe stimuli at the beginning of each trial (**Figure 1A**).

**Figure 5A** shows the responses of an example OFC neuron with these effects. This neuron responded weakly to options as they appeared in sequence, but responded strongly when the correct option appeared. To assess this response statistically, we calculated the average firing rate during each of the three stimulus presentation epochs on all correct trials. We then used two-way t-tests to compare the average firing rate during epochs in which the correct stimulus was presented to the average firing rate during epochs in which the correct stimulus was not presented, separately for each of the three presentation epochs. This cell demonstrated a significantly greater firing rate when the correct stimulus was presented compared to when the correct stimulus was not presented during all three presentation epochs (first epoch: *P* < 0.0001, second epoch: *P* = 0.0048, third epoch: *P* < 0.0001, two-way t-tests).

**Figure 5.**
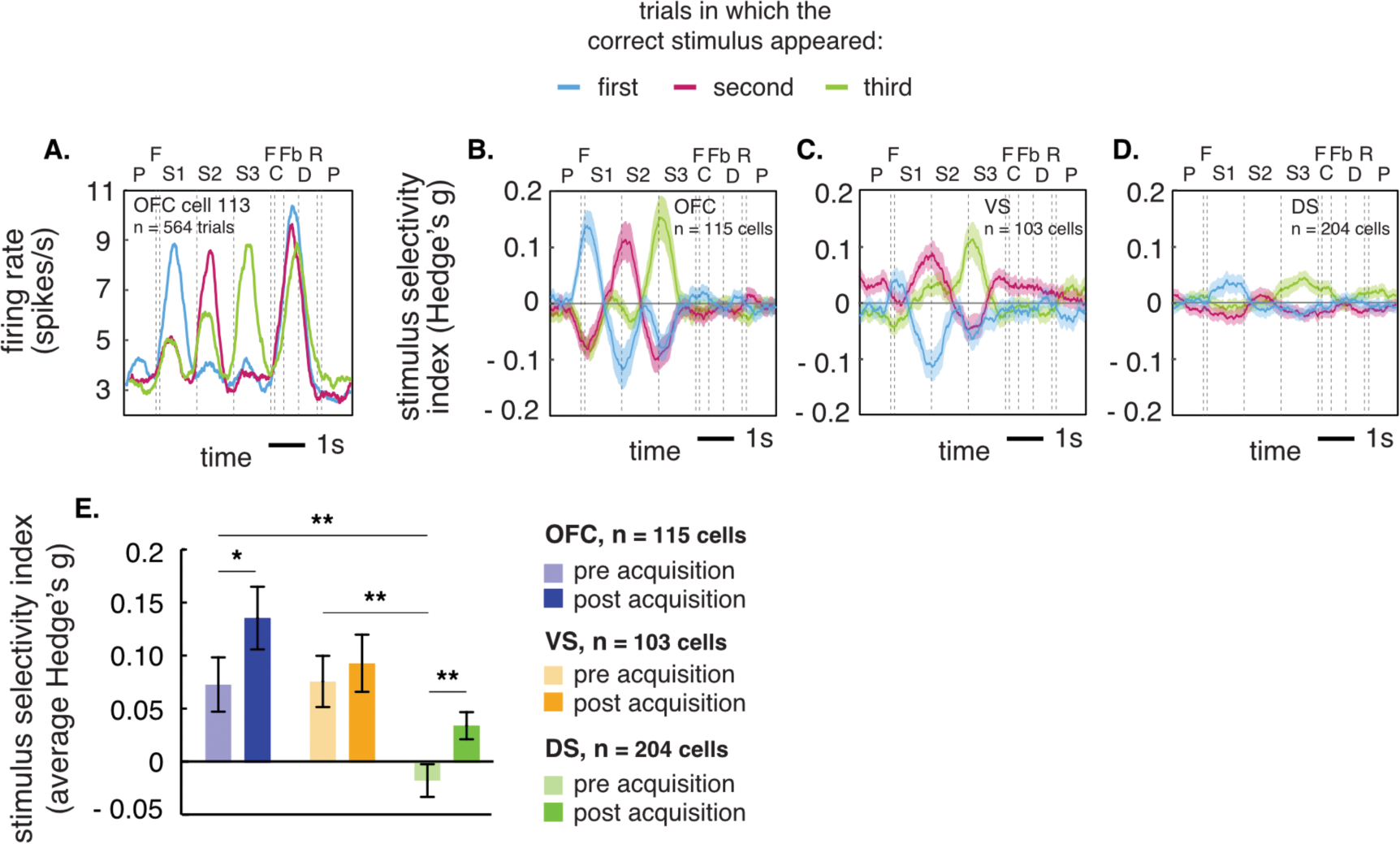
Correct stimulus selectivity. (A) Average response of a single OFC neuron demonstrating selectivity for the presentation of the correct stimulus during the first (blue line), second (red line), and third (green line) presentation epochs. C, choice; Fb, feedback; D, delay; R, reward; P, preparatory period (ITI); F, fixation; S1, first stimulus appearance; S2, second stimulus appearance; S3, third stimulus appearance. (B-D) Correct stimulus selectivity (Hedges g) across the populations of OFC (B), VS (C), and DS (D) neurons. (E) Average proportion of variance explained (Hedge’s g) by the presentation of the correct stimulus before late switch points and after late switch points for the populations of OFC (light blue and dark blue bars), VS (light orange and dark orange bars), and DS (light green and dark green bars) neurons. The analysis epoch for each region consists of a 200 ms period surrounding the average time of maximum selectivity. Bar graph shows the average Hedge’s g (± SEM) during these epochs. Selectivity measures reflect averages across all neurons. * P < 0.05, ** P < 0.01.

A significant proportion of cells in OFC demonstrated modulation associated with the presentation of the correct stimulus during all three presentation epochs (first epoch: 35.65%, n = 41/115 cells; second epoch: 46.96%, n = 54/115 cells; third epoch: 49.57%, n = 57/115 cells, *P* < 0.0001 for all comparisons, binomial tests). Significant percentages were also observed in VS (first epoch: 45.63%, n = 47/103 cells; second epoch: 47.57%, n = 49/103 cells; third epoch: 50.49%, n = 52/103 cells, *P* < 0.0001 for all comparisons) and DS (first epoch: 35.47%, n = 72/204 cells; second epoch: 36.95%, n = 75/204 cells; third epoch: 41.87%, n = 85/204 cells, *P* < 0.0001 for all comparisons).

### Associative learning related activity increases after rule acquisition in OFC and DS, but arises early in learning and remains constant across the block in VS

We next examined the average magnitude of correct-stimulus selectivity using Hedge’s g (a bi-directional effect size measure similar to Cohen’s d, see methods). The average selectivity across time within trials for the population of OFC cells is shown in **Figure 5B**. Data for VS and DS are shown as well, for comparison (**Figure 5C and 5D**). Within trials, we observed that the timing of neural responses in OFC and VS appeared to arise sooner after the presentation of stimuli compared to DS, which is consistent to the general pattern we observed for single neurons. Thus, to directly assess the timing of correct stimulus selectivity, we first determined the average time of maximum selectivity within trials in each region (averaged across all correct trials and all three presentation epochs). We found that correct stimulus selectivity peaked 370 ms after the start of the stimulus presentation period in OFC, 340 ms after the start of the stimulus presentation period in VS, and 520 ms after the start of the stimulus presentation period in DS. To determine whether these latencies were significantly different across populations of neurons, we calculated the time of maximum selectivity across neurons and performed a one-way ANOVA using the factor brain region (OFC, VS, DS). This analysis revealed a significant effect of brain region *(P* = 0.0478), which was due to a significantly greater latency across the population of DS neurons compared to the populations of OFC neurons *(P* = 0.0442, Fisher’s LSD Test) and VS neurons *(P* = 0.0466, Fisher’s LSD Test).

We then examined correct stimulus selectivity before and after late switch points (**Figure 5E**). Because the populations of OFC, VS, and DS neurons demonstrated significantly different latencies for correct stimulus selectivity, we calculated the average selectivity in a 200 ms window surrounding the average time of maximum selectivity for each population of neurons. We calculated this measure for all correct trials before late switch points and all correct trials after late switch points, averaged across all three presentation epochs. We found no difference in the magnitude of selectivity before or after late switch points in the VS *(P* = 0.5145, Fisher’s LSD Test), but found a significantly greater magnitude of selectivity after late switch points compared to before late switch points in OFC (*P* = 0.0171, Fisher’s LSD Test) and DS *(P* = 0.0072, Fisher’s LSD Test). We also found a significantly greater magnitude of selectivity in OFC compared to DS prior to late switch points (*P* = 0.0016, Fisher’s LSD Test) and after late switch points *(P* = 0.0005, Fisher’s LSD Test), and a significantly greater magnitude of selectivity in VS compared to DS prior to late switch points *(P* = 0.0014, Fisher’s LSD Test) and after late switch points (*P* = 0.0431, Fisher’s LSD Test). We found no difference between OFC and VS at either point (before late switch points: *P* = 0.9312, Fisher’s LSD Test, after late switch points: *P* = 0.1982, Fisher’s LSD Test).

Taken together, the above results indicate that OFC and VS neurons demonstrate greater correct stimulus selectivity than DS neurons both before and after rule acquisition, while neurons in both OFC and DS increase selectivity after rule acquisition. These findings are consistent with our results regarding switch modulation. Specifically, the populations of OFC and DS neurons both demonstrate greater switch modulation at the point of rule acquisition compared to early periods of trial and error learning, while both regions also demonstrate an increase in correct stimulus selectivity after rule acquisition.

## DISCUSSION

In the current study, we describe two new findings based on responses of OFC neurons in a version of the WCST. First, we show that OFC neurons demonstrate switch-related modulation. That is, their firing rates change systematically on trials when monkeys adjust strategies. These signals were observed both on early switches, when monkeys abandoned their earlier strategy and, more strongly, on late switches, when monkeys committed to a new strategy. We also observed associative learning signals in OFC neurons. In other words, we found phasic changes in firing rate associated with the presentation of the correct option in a sequence of stimuli, which presumably reflect the learned association between the stimulus and the reward it predicts because of the rule that is used. These putative associative signals were stronger in OFC following rule acquisition; this finding echoes our finding that switch signals in OFC are greater at the point of rule acquisition than at early switch points.

While the OFC is sometimes thought of as a purely economic structure, a great deal of research indicates that it may have executive roles as well; these roles include rule encoding, working memory for both gustatory and abstract information, conflict-monitoring, information-seeking and curiosity, and linking outcomes with information about the spatial world (Blanchard et al., 2015; Kidd and Hayden, 2015; Lara et al., 2009; Luk and Wallis, 2013; Mansouri et al., 2014; Strait et al., 2016; Tsujimoto et al., 2009; Wallis and Miller, 2003; Wallis et al., 2001). Our new findings fit in with these ideas. Taken together, this work suggests that OFC plays an executive role that is complementary to that of executive dorsal structures, such as the dorsolateral prefrontal cortex and the dorsal anterior cingulate cortex. We suspect that, while there are differences between these regions, the differences may not be as simple as economic vs. executive. Instead, they may have to do with variables like informational modality (Lara et al., 2009).

Although several studies suggest that OFC does not contribute to rule-based switching (Dias et al., 1996a, 1996b, 1997), one recent study suggests it may. Specifically, a recent study in rodents indicates that OFC lesions disrupt switching performance (Chase et al., 2012). Our current findings provide confirmatory evidence for this idea and extend upon it in several ways. First, we show that switching is observed at the single unit level, and that switching correlates are observed for both early and late switches. Second, we show that the results are not limited to rodents. Third, our finding that the strength of associative learning signals in OFC increases following late switch signals further suggests that switch signals in OFC may play a role in guiding or initiating stable target identification and selection. Finally, our finding of switch signals in OFC provides a neural basis for a theory heretofore based solely on behavioral patterns following lesions.

These results complement our recent recordings in DS and VS in the same task. In our previous study (Sleezer and Hayden, 2016), we found correlates of switching in both of these regions that resembles those reported here for OFC. We also found correlates of associative encoding as well. One striking finding is the broad similarity across the regions. In another study, we also found a similarity in rule encoding in all three regions (Sleezer et al., 2016). This similarity is reminiscent of a different study using a different task showing functional overlap between OFC and VS in decision processes related to risky choice (Strait et al., 2016). These results endorse the idea that striatum and its cortical inputs can, in many cases, have some overlap in their functions.

This is not to say that OFC and striatum were strictly identical, even when faced with the same task. For example, we previously found that VS neurons demonstrate context-dependent switch signals. In contrast, we did not find context-dependent switch signals in OFC the present study. In addition, while general switch signals appear to be stronger in VS when monkeys switch away from previously relevant rules, these signals in OFC and DS are stronger when monkeys switch to newly relevant rules. These findings suggest that VS may play a greater role in guiding the identification of newly relevant rules when the correct rule is uncertain, while OFC and DS may play a greater role in guiding stable rule selection once the correct rule is known.

The present results complement earlier research from several labs showing task-switching signals in many brain regions, including OFC, striatum, parietal cortex, dorsal anterior cingulate cortex, and even posterior cingulate cortex (Blanchard and Hayden, 2014; Hayden et al., 2010, 2011; Heilbronner and Hayden, 2016; Kamigaki et al., 2009; Mansouri et al., 2006; Sleezer and Hayden, 2016). Taken together, this body of work supports the idea that task-switching is both widespread and distributed, and provides evidence against the idea that this function is the exclusive domain of a small and highly specialized piece of brain tissue.

The associative encoding signals we found were manifest as an enhanced or suppressed response to cues that matched the learned rule. This finding is intriguing because it is the same type of modulation that has previously been linked to target selection. Specifically, neurons in prefrontal and association cortex show significantly enhanced or suppressed responses to to-be-chosen cues when they appear in a sequence of options (Chelazzi et al., 1998; Hayden and Gallant, 2013; Lui and Pasternak, 2011; Mazer and Gallant, 2003). Indeed, what we call rule here would, in such tasks, be called feature-based attention.

The data do not identify the mechanisms by which neurons gain the ability to discriminate the different offers and respond differently to the one that matches the current rule. However, the fact that rule encoding and switching are observed in the same set of neurons that participate in associative encoding raise an interesting possibility. Perhaps the processes associated with learning the new rule cause a change in the response properties of the OFC neurons. This change in responsiveness is observable in the form of tonic changes in firing rate, and these changes are what we call rule encoding (Sleezer et al., 2016). It is then further observable in the form of its direct effect: changes in the responses of the neurons to the offers. This idea is borrowed from the literature on memory-guided decision-making, and is consistent with the idea that rule-based and memory-based decisions reflect common underlying mechanisms (Chelazzi et al., 1998; Hayden and Gallant, 2013; Lui and Pasternak, 2011; Machens et al., 2005).

## Acknowledgements

This research was supported by a R01 (DA038106), and a Brain and Behavior Research Foundation NARSAD award to BYH, and by a NIH Training Fellowship (T32-EY007125) to BJS. We thank Meghan Castagno and Tommy Blanchard for assistance in data collection and analysis and Marc Mancarella for general lab assistance.

## Author contributions

BJS lead the design of the task and designed and performed all analyses. BJS and BYH wrote the manuscript. GL and MDC supervised the data collection.

## Data accessibility statement

All data will be posted on Figshare and data and code will be available at haydenlab.com/data.

